# Complementary roles for ventral pallidum cell types and their projections in relapse

**DOI:** 10.1101/533554

**Authors:** Asheeta A. Prasad, Caroline Xie, Chanchanok Chaichim, Simon Killcross, John M. Power, Gavan P. McNally

**Author notes:** Correspondence to: Gavan P. McNally PhD School of Psychology UNSW Sydney, NSW Australia.

## Abstract

Ventral pallidum (VP) is a key node in the neural circuits controlling relapse to drug seeking but how this role relates to different VP cell types and their projections is poorly understood. Using male rats, we show how different forms of relapse to alcohol-seeking are assembled from VP cell types and their projections to lateral hypothalamus (LH) and ventral tegmental area (VTA). First, we used RNAScope in situ hybridization to characterize activity of different VP cell types during relapse to alcohol-seeking provoked by renewal (context-induced reinstatement). We found that VP Gad1 and parvalbumin (PV), but not vGlut2, neurons show relapse-associated changes in c-Fos expression. Next, we used retrograde tracing, chemogenetic, optogenetic, and electrophysiological approaches to study the roles of VP^Gad1^ and VP^PV^ neurons in relapse. We show that VPGad1 neurons contribute to contextual control over relapse (renewal), but not to relapse during reacquisition, via projections to LH where they converge with ventral striatal inputs onto LH^Gad1^ neurons. This convergence of striatopallidal inputs at the level of individual LH^Gad1^ neurons may be critical to balancing propensity for relapse versus abstinence. In contrast, VP^PV^ neurons contribute to relapse during both renewal and reacquisition via projections to VTA but not LH. These findings show complementary roles for different VP cell types and their projections in relapse. VP^Gad1^ neurons control relapse during renewal via projections to LH whereas VP^PV^ neurons control relapse during both renewal and reacquisition via projections to VTA. Targeting these different pathways may provide tailored interventions for different forms of relapse.

## Introduction

Drug use contributes significantly to the burden of disease (Whiteford et al., 2013). A treatment goal is promoting abstinence but a consistent propensity to relapse impedes this (Jonas et al., 2014). The ventral pallidum (VP) is a key brain region controlling relapse. VP, together with ventral striatum, forms the ventral striatopallidal system (Heimer et al., 1982; de Olmos and Heimer, 1999). Although initially viewed as controlling motor output (Mogenson et al., 1980), the ventral striatopallidal system has broad functions in motivation including evaluating hedonic impact of rewards and mediating the effects of reward predictive cues on decision making (Smith, 2005; Smith et al., 2009; Ho and Berridge, 2013; Leung and Balleine, 2013; Castro et al., 2015; Leung and Balleine, 2015; Root et al., 2015; Richard et al., 2016). Lesion and pharmacological (McFarland and Kalivas, 2001; McFarland et al., 2004; Tang et al., 2005; Perry and McNally, 2013), circuit specific (Stefanik et al., 2013; Heinsbroek et al., 2017), as well as human lesion (Moussawi et al., 2016) studies identify VP as critical for relapse to drug seeking. This role is shared across different forms of relapse (cue, stress-, context-, and drug-priming reinstatement) to seeking a variety of drugs of abuse (alcohol, psychostimulants, opiates). Hence, VP is an obligatory node in a final common neural circuit for relapse to drug seeking (Kalivas and O'Brien, 2007).

However, the heterogeneity of VP hinders understanding its role in relapse. VP comprises a variety of different cell types (glutamatergic, GABAergic, and cholinergic) that express different neuropeptides (Root et al., 2015) with diverse electrophysiological characteristics (Napier and Mitrovic, 1999; Root et al., 2010; Kupchik and Kalivas, 2012; Avila and Lin, 2014). Further, VP neurons form multiple output pathways, including to lateral hypothalamus, subthalamic nucleus, lateral habenula, and ventral tegmental area as well as prefrontal cortex, amygdala, and nucleus accumbens (Haber et al., 1985; Zahm et al., 1987; Root et al., 2015). Recent work has begun to show how this cell-type and projection heterogeneity relates to different behavioral states (Knowland et al., 2017; Faget et al., 2018; Tooley et al., 2018). Yet, how the role for VP in relapse relates to this cell-type and projection heterogeneity remains poorly understood. Here we used an animal model of relapse to address this.

Across nine experiments we used renewal (context-induced reinstatement) (Crombag and Shaham, 2002; Crombag et al., 2008; Khoo et al., 2017) and reacquisition (Willcocks and McNally, 2011) models of relapse to alcohol-seeking to study the roles of three major VP cell populations – Gad1, Parvalbumin (PV), and vGlut2 – in relapse. First, we used RNAScope *in situ* hybridization for c-Fos and Gad1, PV, or vGlut2 mRNA to assess recruitment of VP cell populations during renewal. Next, we used retrograde tracing, chemogenetic, optogenetic, and electrophysiological approaches to study the causal roles of VP^Gad1^ and VP^PV^ neurons and their projections to the lateral hypothalamus and ventral tegmental area in relapse. We show that VP^Gad1^ neurons control relapse during renewal via projections to LH whereas VP^PV^ neurons control relapse during both renewal and reacquisition via projections to VTA.

## Materials and Methods

### Subjects

Subjects were adult male Sprague-Dawley (Animal Resources Centre; Perth, Australia), LE-Tg(Gad1-iCre)3Ottc and LE-Tg(PV-iCre)3Ottc (Optogenetics and Transgenic Technology Core, NIDA, MD, USA obtained via Rat Research Resource Centre RRRC#751, RRRC#773). They were housed in ventilated racks, in groups of 4, on corn cob bedding in a climate-controlled colony room maintained on 12:12 hour light/dark cycle (0700 lights on). Rats in behavioral studies had free access to food (Gordon’s Rat Chow) and water until two days prior to commencement of behavioral training when they received 1 hour of access to food and water each day for the remainder of the experiment. All subjects were randomly allocated to experimental conditions. All studies were performed in accordance with the Animal Research Act 1985 (NSW), under the guidelines of the National Health and Medical Research Council Code for the Care and Use of Animals for Scientific Purposes in Australia (2013). The UNSW Animal Care and Ethics Committee approved all procedures.

### Surgeries and injections

Rats were anaesthetized via i.p. injection with a mixture of 1.3 ml/kg ketamine anaesthetic (Ketamil; Troy Laboratories, NSW, Australia) at a concentration of 100 mg/ml and 0.3 ml/kg of the muscle relaxant xylazine (Xylazil; Troy Laboratories, NSW, Australia) at a concentration of 20 mg/ml. Rats received a subcutaneous injection of 0.1ml 50mg/ml carprofen (Pfizer, Inc., Tadworth, United Kingdom) before being placed in the stereotaxic frame (Kopf Instruments, CA, USA). They received stereotaxic surgery using the following flat skull coordinates relative to bregma (A-P, M-L, D-V in mm): VP AAV +0.00, ±2.50, −8.50; LH AAV and tracing −2.60, ±1.80, −8.60; LH optic fibres −2.60, ±3.60, −8.80 (10° angle); AcbSh AAV +1.28, ±0.70, −8.20; VTA AAV −5.80, ±2.25, −8.30 (10° angle).

Vectors and tracers were infused with a 23-gauge, cone tipped 5μl stainless steel injector (SGE Analytical Science, Australia) over 3 min using an infusion pump (UMP3 with SYS4 Micro-controller, World Precision Instruments, Inc., FL, USA). The needle was left in place for 7 min to allow for diffusion and reduce spread up the infusion tract. Optogenetic cannulae for relevant experiments were implanted during a concurrent stereotaxic procedure. Hand fabricated cannula were secured using jeweller’s screws and dental cement (Vertex Dental, The Netherlands). At the end of surgery, rats received intramuscular injection of 0.2 ml of 150 mg/ml solution of procaine penicillin (Benacillin; Troy Laboratories, NSW, Australia) and 0.2 ml of 100 mg/ml cephazolin sodium (AFT Pharmaceuticals, NSW, Australia). All rats were monitored daily for weight and/or behavioral changes.

### Viral vectors

pAAV-hSyn-DIO-hM4D(Gi)-mCherry was a gift from Bryan Roth (Addgene viral prep # 44362-AAV5; http://n2t.net/ addgene:44362; RRID:Addgene_44362, 5.4×10^e12^ vp/mL). pAAV-hSyn-hM4D(Gi)-mCherry was a gift from Bryan Roth (Addgene viral prep # 50475-AAV5; http://n2t.net/addgene:50475; RRID:Addgene_50475, 4.1×10^e12^ vp/ml). pAAV-hSyn-eNpHR 3.0-EYFP was a gift from Karl Deisseroth (Addgene viral prep # 26972-AAV5; http://n2t.net/addgene:26972; RRID:Addgene26972, 7.7×10^e12^ vp/ml). pAAV-Syn-ChrimsonR-tdT was a gift from Edward Boyden (Addgene viral prep # 59171-AAV5; http://n2t.net/addgene:59171; RRID:Addgene59171, 2.9×10^e12^ vp/mL). pAAV-EF1a-double floxed-hChR2(H134R)-EYFP-WPRE-HGHpA was a gift from Karl Deisseroth (Addgene viral prep # 20298-AAV5; http://n2t.net/addgene:20298; RRID:Addgene_20298, 7.4×10^e12^ vp/ml). AAV5-hSyn-DIO-ChR2(H134R)-eYFP AAV5-hSyn-eYFP (4.9×10^e12^ vp/ml) and AAV5-hSyn-DIO-eYFP (4.1×10^e12^ vp/mL) were obtained from UNC Vector core from plasmids as gifts from Karl Deisseroth.

### Apparatus

Standard rat operant chambers (ENV-008) (Med Associates, VT, USA) were used for all behavioral procedures. The chambers contained two nosepoke holes symmetrically located on one sidewall of the chamber, 3cm above a grid floor. A recessed magazine was located behind a 4×4cm opening in the center of the same wall between the two nosepokes. Responding on one (active) nosepoke extinguished the cue light in the nosepoke and triggered a syringe pump to deliver alcoholic beer to the magazine during acquisition training, whereas responding on the other (inactive) nosepoke had no programmed consequences. A computer running MedPC-IV software (Med Associates, VT, USA) controlled all events. Suspended above each chamber was an LED plus fibre-optic rotary joint and LED driver (Doric Lenses Inc., Quebec, Canada) controlled by MedPC-IV. The eight self-administration chambers were divided into two groups of four to serve as distinct contexts for experiments with context as a factor. These chambers differed in their olfactory (rose vs peppermint essence), tactile (grid vs Perspex flooring) and visual (light on vs off) properties. These two contexts were fully counterbalanced to serve as the training (context A) and extinction (context B) contexts. Fibre optic cannulae and patch cables were fabricated from 0.39 NA, Ø400μm core multimode optical fiber and ceramic ferrules (Thor Labs, Newton, NJ, USA). Plexiglas chambers (Med Associates, VT, USA) with dimensions 43.2cm (width) × 43.2cm (length) × 30.5cm (height) were used for locomotor assessment. Movement was tracked with three 16 beam infrared arrays. Infrared beams were located on both the x- and y-axes for positional tracking.

### Behavioral Procedures

#### General behavioral testing procedures

On the first two days, the animals received 20 min magazine training sessions in both context A and context B. During these sessions, there were 10 non-contingent deliveries of 0.6ml of the reward (4 % alcohol (v/v) decarbonated beer; Coopers Brewing Company, Australia) at time intervals variable around a mean of 1.2 min. On the next ten days rats received self-administration training in context A for 1 hr per day. Responding on the active nosepoke extinguished the nosepoke cue light and triggered delivery via syringe pump of 0.6 ml alcoholic beer to the magazine on an FR-1 schedule followed by a 24 s timeout. Responses on the inactive nosepoke were recorded but had no consequences. On the next four days, rats received extinction training in context B for 1 h per day. During this training, responses on the active nosepoke extinguished the cue light and triggered the pump but no beer was delivered. All rats were tested starting 24 hours after the last extinction session. We selected these procedures based on our past work that has shown robust context-induced reinstatement and reacquisition under these conditions (Hamlin et al., 2006; 2007; 2008; Gibson et al., 2018).

For renewal, rats were tested twice, once in the extinction context (extinction, context B) and once in the training context (reinstatement, context A). Tests lasted 30 (optogenetic experiments) or 60 min (chemogenetic experiments). The order of tests was counterbalanced and tests were identical to self-administration except that the syringe pump was empty. 24 hr after extinction/reinstatement test, rats were tested for the reacquisition of alcohol-seeking in a single session in the training context. Tests lasted 30 (optogenetic experiments) or 60 min (chemogenetic experiments). For chemogenetic manipulation, rats received an i.p. injection 0.1 mg/kg of clozapine (#C6305, Sigma Aldrich; 1 ml/kg diluted, 5% DMSO and saline) 30 min prior to tests. For optogenetic manipulation, rats were connected to patch cables attached to 625nm LEDs (Doric Lenses Inc., Quebec, Canada). 625nm (constant, 8-10mW) light was delivered upon commencement of the test sessions.

#### Profile of VP subpopulations during renewal

There were three groups; Groups AB0 (n = 4), ABB (n = 6) and ABA (n = 6), All rats were trained to respond for alcoholic beer in context A and extinguished in context B. Rats were tested once, for 1 hr, 24 hr after the last extinction session. Group ABB was tested in the extinction context. Group ABA was tested in the training context. Group AB0 was not tested, instead they were simply transported to the laboratory on test and received an equivalent period of handling. This group served as a control for the behavioral and pharmacological training histories. All rats were sacrificed 30 min after test (or at the equivalent time for Group AB0). Their brains were later processed for expression of the cFos mRNA with GAD1, vGlut2 or PV using RNAscope technology. Images from 3 sets from the VP were obtained from each brain; rostral, medial and caudal VP. Images were analysed for single-labelled and double-labelled cells using Adobe Photoshop (Adobe Systems, CA, USA).

#### Chemogenetic inhibition of VP^GAD1^ neurons on renewal, reacquisition and locomotor activity

There were two groups, VP^GAD1^ Cre− (n = 8) and VP^GAD1^ Cre+ (n = 7). Both groups received AAV5-hSyn-DIO-hM4D(Gi)-mCherry, bilaterally in the VP. Rats were trained and extinguished as described above. Testing commenced 24 hr after extinction. Rats were tested for 1 hr in the extinction context (ABB) and for 1 hr in the training context (ABA) for expression of extinction and renewal (context-induced reinstatement), respectively. These tests were separated by 24 hr and the order of tests was counterbalanced within-subjects. Rats were tested 24 hr later for 1 hr reacquisition of alcoholic beer seeking in the training context. Our past research has shown no impact of the prior order of ABA and ABB testing on responding during reacquisition.

#### Chemogenetic inhibition of VP^PV^ neurons on renewal, reacquisition and locomotor activity

There were two groups, VP^PV^ Cre− (n = 8) and VP^PV^ Cre+ (n = 7). Both groups received AAV5-hSyn-DIO-hM4D(Gi)-mCherry. All rats were then trained, extinguished, and tested for expression of extinction (ABB), renewal (ABA), reacquisition and locomotor test as described in the general behavioral testing procedures.

#### VP output pathways recruited during renewal of alcohol seeking

To assess recruitment of the VP→LH pathway during context-induced reinstatement we applied the retrograde tracers CTb-488 (Thermofisher Scientific, Cat# C34775) and CTb-555 (Thermofisher Scientific, Cat# C34776) (counterbalanced) into LH. There were three groups: Group ABB (n = 6) (tested in the extinction context), group ABA (n = 5) (tested in the training context), and group AB0 (n = 3) (transported to the laboratory on test, handled, but not tested). The behavioral procedure was the same as above, except all rats were perfused 1 hr after the 1 hr test (or at the equivalent time for Group AB0). Their brains were later processed for immunohistochemistry for c-Fos protein and CTb tracer.

#### Optogenetic disconnection of VP→LH pathway on renewal and reacquisition

We used AAV5-hSyn-eYFP and AAV5-hSyn-eNpHr3.0-eYFP bilaterally infused into the VP and optic cannula bilaterally implanted above the LH. A minimum of 4 weeks after surgery, eYFP (n = 8) or eNpHR3.0 (n = 6) groups received training, extinction, tested for expression of extinction (ABB), renewal (ABA), and reacquisition as described in the general behavioral testing procedures.

#### Chemogenetic disconnection of VP^GAD1^-LH pathway on renewal and reacquisition and locomotor activity

There were two groups, VP^GAD1^ Cre− (n = 5) and VP^*GAD1*^ Cre+ (n = 5). Both groups received AAV5-hSyn-DIO-hM4D(Gi)-mCherry in one VP and AAV5-hSyn-hM4D(Gi)-mCherry in the contralateral LH. Hemispheres were counterbalanced across animals. All rats were trained, extinguished, and tested for expression of extinction (ABB), renewal (ABA), reacquisition and locomotor test as described in the general behavioral testing procedures.

#### Chemogenetic disconnection of VP^PV^ − LH on renewal, reacquisition and locomotor activity

There were three groups, VP^PV^ Cre− (n = 6) contralateral disconnection, VP^PV^ Cre+ (n = 7) ipsilateral disconnection and VP^PV^ Cre+ (n = 7) contralateral disconnection. All groups received AAV5-hSyn-DIO-hM4D(Gi)-mCherry in the VP and AAV5-hSyn-hM4D(Gi)-mCherry in the contralateral or ipsilateral LH, counterbalanced across animals. All rats were trained, extinguished, and tested for expression of extinction (ABB), renewal (ABA), reacquisition and locomotor activity as described in the general behavioral testing procedures.

#### Chemogenetic disconnection of VP^PV^ − VTA on renewal, reacquisition and locomotor activity

There were three groups, VP^PV^ Cre− (n = 7) contralateral disconnection, VP^PV^ Cre+ (n = 6) ipsilateral disconnection and VP^PV^ Cre+ (n = 9) contralateral disconnection. All groups received AAV5-hSyn-DIO-hM4D(Gi)-mCherry in the VP and AAV5-hSyn-hM4D(Gi)-mCherry in the ipsilateral or contralateral VTA. All rats were trained, extinguished, and tested for expression of extinction (ABB), renewal (ABA), reacquisition and locomotor activity as described in the general behavioral testing procedures.

### Electrophysiology

Brain slices were prepared from Gad1-cre+ rats that had received AAV5-DIO-eYFP to LH, AAV5-hSyn-Chrimson-tdTomato to AcbSh, and AAV5-DIO-hChR2(H134R)-eYFP to VP at least 6 weeks prior to slice preparation. Rats were deeply anaesthetized with Isoflurane (5%), decapitated and their brain rapidly removed and submerged in ice-cold oxygenated (95% O2, 5% CO2) HEPES based artificial cerebral spinal fluid (HEPES-aCSF; 95 mM NaCl, 2.5 mM KCl, 30 mM NaHCO_3_, 1.2 mM NaH_2_PO_4_, 20 mM HEPES, 25 mM glucose, 5 mM ascorbate, 2 mM thiourea, 3 mM sodium pyruvate, pH adjusted to 7.3 − 7.4 in NaOH) with low (0.5 mM) CaCl_2_, and high (10 mM) MgSO_4_ for 2 − 3 min. Coronal slices (300 μm) were prepared using a vibratome (model VT1200, Leica, Wetzlar, Germany), incubated for 10 min in a 30°C in a neural protective recovery HEPES-aCSF (NaCl was replaced by equimolar N-Methyl-D-glucamine; pH adjusted to 7.3-7.4 with HCl) and then transferred to a Braincubator (#BR26021976, Payo Scientific, Sydney, Australia) and maintained at 16°C in a HEPES-aCSF holding solution with 2 mM CaCl_2_, and 2 mM MgSO_4_.

For recordings, slices were transferred to a recording chamber and continuously perfused (2-3 ml min-1) with aCSF (30°C) containing: 119 mM NaCl, 5 mM KCl, 1.3 mM MgCl_2_, 2.5 mM, CaCl_2_, 1.0 mM NaH_2_PO_4_, 26.2 mM NaHCO_3_, 11 mM glucose (equilibrated with 95% CO_2_, 5% O_2_). Targeted whole-cell patch clamp recordings were made from visually identified neurons using a microscope (Zeiss Axio Examiner D1) equipped with a 20x water immersion objective (1.0 NA) a LED fluorescence illumination system (pE-2, CoolLED, Andover, UK) and an EMCCD camera (iXon+, Andor Technology, Belfast, UK). Patch pipettes (2 – 5 MΩ) were filled with an internal solution containing (in mM) 130 potassium gluconate, 10 KCl, 10 HEPES, 4 Mg_2_-ATP, 0.3 Na_3_-GTP, 0.3 EGTA, 10 phosphocreatine disodium salt (pH 7.3 with KOH, 280 − 290 mOsm). Electrophysiological recordings were amplified using a Multiclamp 700B amplifier (Molecular Devices, California, USA), filtered at 6 kHz, digitized at 20 kHz with a Digidata1440A (Molecular Devices) interface, and controlled using AxoGraph (Axograph, Sydney, Australia). Electrophysiological data were analyzed offline using Axograph.

Optogenetic stimulation of ChR2 and Chrimson expressing neurons was elicited using 470 nm or GYR (excitation filter 605 nm, 50 nm FWHM) LED illumination delivered through the objective. Neurons were considered connected if the light reliably evoked a short latency synaptic current > 5 pA (Vm = − 50 mV). Light-evoked responses were averaged. The latency of light-evoked responses was calculated from light onset. Picrotoxin (Tocris) was dissolved in DMSO, aliquoted and stored frozen (−30°C) as concentrated (1000 – 4000x) stock solution until needed.

### Histology and Immunohistochemistry

#### RNAscope in situ hybridization

*In situ* hybridization was performed as previously described (Rubio et al., 2015; Gibson et al., 2018). Thirty minutes post-test session, rats were deeply anesthetized with sodium pentobarbital (100 mg/kg, i.p.; Virbac, NSW, Australia) and decapitated. Brains were rapidly extracted and quick frozen on quick freeze (Lecia Biosystems, NSW, Australia). Brains were stored at 80°C until use. 16 μm coronal sections of VP were cut using a cryostat and mounted directly onto Super Frost Plus slides (Fisher Scientific, NH, USA). Slides were stored at −80 °C until use. The RNAscope Multiplex Fluorescent Reagent Kit (Advanced Cell Diagnostics, #320850) was used to detect GAD1 (316401-C2, GenBank ID NM_017007.1), vGlut2 (317011-C2, GenBank ID; NM_053427.1), PV (407821-C3, GenBank ID; NM_022499.2) and cFos (403591,GenBank ID; NM_022197.2) mRNA in VP. Slides were fixed in 4% paraformaldehyde for 15mins at 4C. Slides were then rinsed in PBS followed by dehydration in increasing concentrations of ethanol (50%, 70%, 100% and 100%) for 10 mins at room temperature then air-dried for 10 mins and a hydrophobic barrier was drawn around the brain sections. Sections were then protease treated (pre-treatment 4) at room temperature for 15 min. Next, sections were then rinsed in distilled water followed by probe application. The target probes used were GAD1 and cFos, vGlut2 and cFos, PV and cFos. Slides were incubated with probes at 40°C for 1 hr. Following this, slides were incubated with preamplifier and amplifier probes (AMP1, 40°C for 30min; AMP2, 40°C for 15min; AMP3, 40°C for 30min). Slides were incubated in fluorescently labelled probes by selecting a specific combination of colors associated with each channel i.e., GAD1 and c-Fos, vGlut2 and c-Fos, PV and cFos. AMP4 Alt B to detect c-Fos/GAD1 and AMP4 Alt C to detect c-Fos/PV in Atto 550 channel (PV or GAD1) and c-Fos in Atto 647 channel. Finally, sections were incubated for 20 s with DAPI and coverslipped with Permafluor mounting medium (Thermo Fisher Scientific, Waltham, Massachusetts). Fluorescent images of the VP were taken with an Olympus BX51 microscope using a 40x objective. VP sections were taken anterior, medial and caudal at Bregma 0.00mm, −0.12, −0.24 (rat brain atlas). Photoshop software was used to quantify the number of neurons containing GAD1, vGlut2, PV, c-Fos and colocalized c-Fos mRNA.

#### Retrograde tracing

The retrograde tracers were cholera toxin b subunit (CTb) conjugated to Alexa Fluor 488 (CTb-488) (Cat#C34775 ThermoFisher Scientific) or Alexa Fluor 555 (CTb-555) (Cat#C34776, ThermoFisher Scientific ThermoFisher Scientific, Australia). Tracers were targeted to the LH. At least one week after surgery, rats were deeply anesthetized with sodium pentobarbital (100 mg/kg, i.p.; Virbac, Milperra, Australia) and perfused transcardially with 200 mL of 0.9% saline, containing heparin (360 mL/L) and sodium nitrite (12.5mL/L), followed by 400 mL of 4% paraformaldehyde in 0.1M phosphate buffer (PB), pH7.4. Brains were extracted from the skull and postfixed for 1 hr in the same fixative and then placed in 20% sucrose solution overnight. Brains were frozen and sectioned coronally at 40 mm using a cryostat (Leica CM1950). Four serially adjacent sets of tissue from the VP and LH from each brain and stored in 0.1% sodium azide in 0.1MPBS, pH 7.2. Brain tissue was processed for c-Fos immunochemistry as described above. Brain slices were mounted onto glass slides, dried for approximately 5 min and coverslipped with Permafluor mounting medium.

#### eYFP/mCherry/cFos immunohistochemistry

In all other experiments, rats were deeply anesthetized with sodium pentobarbital (100 mg/kg, i.p.; Virbac, NSW, Australia) and perfused transcardially with 200 mL of 0.9% saline, containing heparin (360 mL/L) and sodium nitrite (12.5mL/L), followed by 400 mL of 4% paraformaldehyde in 0.1M phosphate buffer (PB), pH7.4. Brains were extracted from the skull and post fixed for 1 hr in the same fixative and then placed in 20% sucrose solution overnight. Brains were frozen and sectioned coronally at 40 mm using a cryostat (Leica CM1950). To visualize eYFP, mCherry immunoreactivity (-IR) (Rabbit anti-eGFP Polyclonal Antibody, Cat#AA11122; RRID AB_221569; Rabbit anti-mCherry Polyclonal Antibody, Cat#PA5-34974; RRID AB_2552323 ThermoFisher Scientific) four serially adjacent sets of sections from the regions of interest were obtained from each brain and stored in 0.1% sodium azide in 0.1M PBS, pH 7.2. Sections were washed in 0.1M PB, followed by 50% ethanol, 50% ethanol with 3% hydrogen peroxidase, then 5% normal horse serum (NHS) in PB (30 min each). Sections were then incubated in rabbit antiserum against eGFP or mCherry (1:2000; ThermoFisher Scientific, MA, USA) in a PB solution containing 2% NHS and 0.2% Triton X-10 (48 hr at 4C). The sections were then washed and incubated in biotinylated donkey anti-rabbit (1:1000; 24 hr at 4C; Biotin Donkey Anti-Rabbit Cat#711-065-152; RRID:AB_2540016 Jackson ImmunoResearch Laboratories, PA, USA). Finally, sections were incubated in avidin-biotinylated horseradish peroxidase complex (6 mL/mL avidin and 6 mL/mL biotin; 2 hr at room temperature; Vector Laboratories, CA, USA, #PK-6100), washed in PB, and then incubated for 15 min in a diaminobenzidine solution (DAB) containing 0.1% 3,3-diaminobenzidine, 0.8% D-glucose and 0.016% ammonium chloride. Immunoreactivity was catalyzed by the addition of 0.2 mL/mL glucose oxidase aspergillus (24 mg/mL, 307 U/mg, Sigma-Aldrich, NSW, Australia). Brain sections were then washed in PB, mounted onto gelatin coated slides, dehydrated, cleared in histolene, and coverslipped with Entellan (Proscitech, Kirwin, Australia), and assessed using an Olympus BX51transmitted light microscope (Olympus, Shinjuku, Tokyo, Japan).

For detection of c-Fos, brain sections were then incubated in rabbit anti-c-Fos (1:500, Santa Cruz Biotechnology Cat# sc-52, RRID:AB_2106783) for 24 h at 4°C. The primary antibodies were diluted in blocking buffer. After washing off unbound primary antibody, sections were incubated overnight at 4°C in biotinylated donkey anti-rabbit IgG (1:500; Jackson ImmunoResearch Labs Cat# 711-065-152, RRID:AB_2340593) diluted in blocking buffer. After washing off unbound secondary antibody, sections were incubated for overnight at 4°C in streptavidin, AlexaFluor-350 conjugate (1:300, S11249, Invitrogen). Brain sections were then washed in PB, pH 7.4, and mounted using mounting media. VP sections were delineated according to Paxinos and Watson (2007) and imaged at 20x using a transmitted light microscope (Olympus BX51). Counts of all neurons immunoreactive (IR) for c-Fos and CTb-488 or CTb-555 native fluorescence across 3 VP sections (80 μm apart) were made using Photoshop (Adobe).

Fluorescent images of VP were taken microscope (Olympus BX51). Counts of all neurons immunoreactive (IR) for c-Fos and CTb-488 or CTb-555 native fluorescence across 3 VP sections (80 μm apart) were made using Photoshop (Adobe).

### Quantification and Statistical Analyses

Data in figures are represented as mean ± SEM unless otherwise stated. Group sizes were based on our past experience with these preparations showing that they were sufficient to detect large (d = 0.8) effect sizes in neuroanatomical or behavioral studies with at least 80% power. The criteria for inclusion in final analysis was correct AAV or tracer and/or fiber placements determined after histology. Group numbers used for analyses in each experiment are indicated at two locations: (1) under the subheadings of behavioral procedures above; (2) in the main results text below. Our primary behavioral dependent variables were numbers of active nosepokes, inactive nosepokes, and distance travelled (locomotor activity). These data were analyzed by means of ANOVA and analyses involving repeated measures adopted a multivariate approach (Boik, 1981; Harris, 2004). All analyses partitioned variances into main effect and interaction terms using Psy Statistical Package (Bird, 2004).

## Results

### Distinct roles for ventral pallidum cell types in relapse

To identify specific VP cell types recruited during relapse, we used ABA renewal of alcohol seeking (Crombag and Shaham, 2002; Bouton and Todd, 2014). Three groups of rats were trained to respond for alcoholic beer in a distinctive context (context A). Responses on one nosepoke (active) led to delivery of alcoholic beer to a magazine cup, whereas responses on a second (inactive) did not. Active nosepoking increased across training. Then we extinguished this behavior in a second context where responses did not earn alcohol (context B). As expected, active nosepoking declined. The mean SEM levels of responding at the end of training and extinction in this and remaining experiments are shown in **Table 1.** In this and remaining experiments, there were no significant differences between groups in nosepokes in self-administration or extinction (all p > .05). One group was tested for expression of extinction (ABB, n = 6), one for relapse (ABA, n = 6), and a third group remained in their home cages and were not tested (AB0, n = 4). This AB0 group served as a control for the behavioral training and pharmacological histories of the groups tested for renewal (Hamlin et al., 2006; 2007; 2008).

Responding was low for group ABB tested in the extinction context. In contrast, responding was higher for group ABA tested in the training context (Context × Nosepoke interaction *F* _(1,10)_ = 7.58, *p* = .020) (**Figure 1A**). This shows renewal of alcohol-seeking (i.e. context-induced reinstatement). All rats were perfused 30 min after test and relapse-associated activity in VP GABA (VP^Gad1^), glutamate (VP^vGlut2^), and parvalbumin (VP^PV^) neurons was assessed via *in situ* hybridization for c-Fos mRNA. ABA renewal was associated with significant c-Fos expression (i.e. ABA > ABB = AB0) in VPGad1 (*F* _(1, 13)_ = 17.00; p = .0012) and VP^PV^ (*F* _(1, 13)_ = 14.61; p < .0021) neurons (**Figure 1B**, **Table 2**). In contrast, VP^vGlut2^ neurons were strongly recruited regardless of test context and their activity, at least as measured by c-Fos mRNA expression, was unrelated to differences in relapse behavior (i.e. ABA = ABB > AB0) (*F* _(1, 13)_ = 27.87; p = .000149).

**Figure 1.**
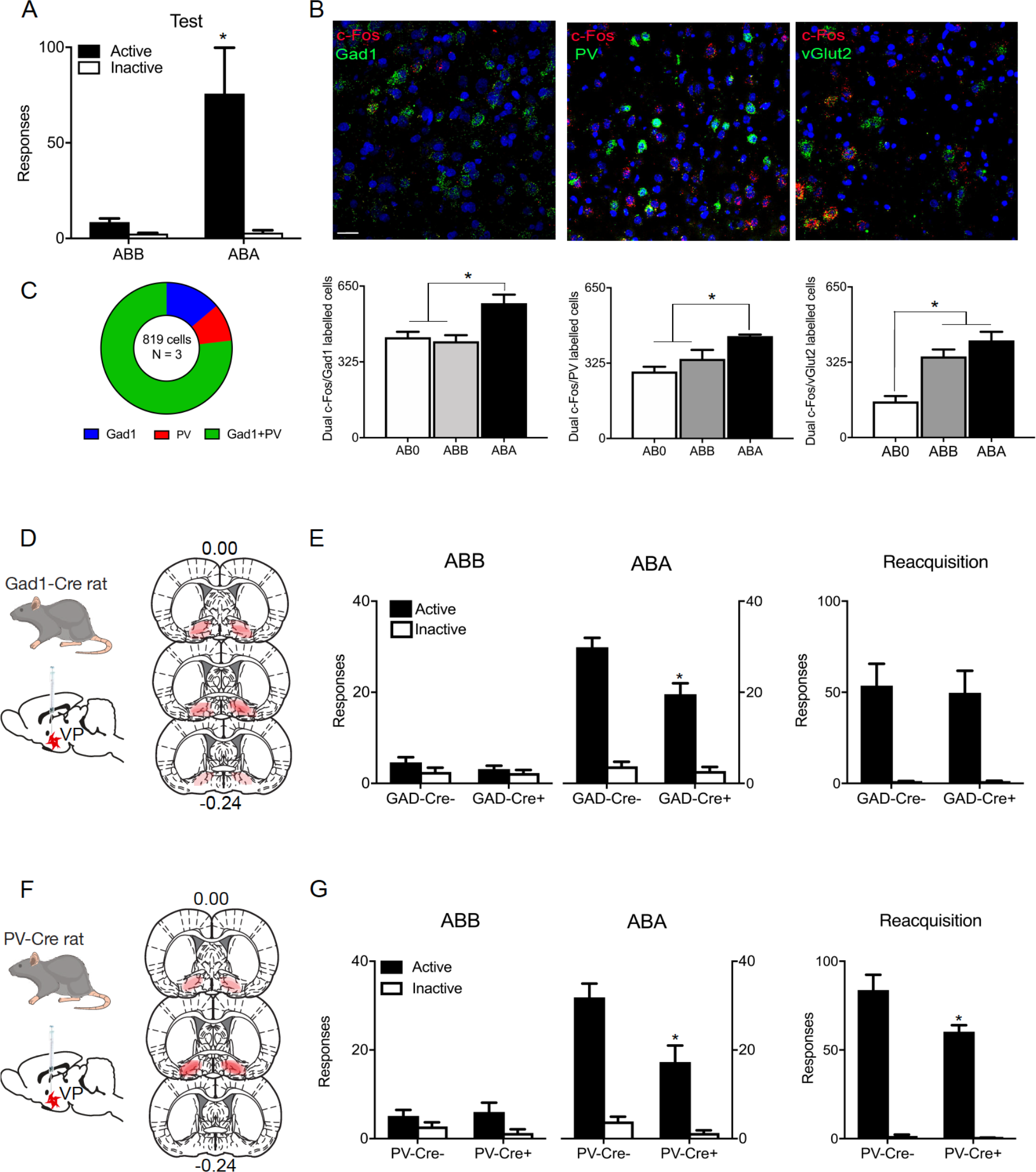
Distinct roles for pallidal cell types in relapse. (A), ABA renewal of extinguished alcohol seeking. (B), ABA renewal is associated with c-Fos mRNA expression in VP Gad1 and PV neurons whereas vGlut2 neurons express c-Fos mRNA independently of test context. Scale bar = 30 m. (C), There was substantial overlap in VP cell populations as shown by co-expression of Gad1 and PV mRNA expression. (D), Cre-dependent inhibitory hM4Di was applied to VP of Gad1-Cre rats, with location of hM4Di expression shown for all animals at 10% opacity. (E), Chemogenetic inhibition of VP Gad1 neurons reduced relapse during renewal but not reacquisition. (F), Cre-dependent inhibitory hM4Di was applied to VP of PV-Cre rats, with location of hM4Di expression shown for all animals at 10% opacity. (G), Chemogenetic inhibition of VP PV neurons reduced relapse during renewal and reacquisition.

The identities of VP neurons overlap (Root et al., 2015), in particular VP^PV^ neurons express vesicular transporters for GABA (vGat) or glutamate (vGlut2) and transmit either inhibitory or excitatory signals (Knowland et al., 2017). The common recruitment of VP^Gad1^ and VP^PV^ neurons during relapse could reflect this overlap. To assess this, we measured PV and Gad1 mRNA in 819 VP neurons from a separate group of animals (n = 3). Gad1 and PV mRNA were co-expressed in a large number of VP neurons with a minority expressing one but not the other (**Figure 1C**).

Next, we determined the roles of VP^Gad1^ and VP^PV^ neurons in relapse. We used adenoassociated viral vectors (AAV) in LE-Tg(Gad1-iCre)3Ottc (Gad1-Cre) rats (*n* = 8) or Gad1-Cre− (*n* = 7) rats (Sharpe et al., 2017) to express the inhibitory hM4Di designer receptor (AAV5-hSyn-DIO-hM4Di-mCherry) (Armbruster et al., 2007) in VP^Gad1^ neurons (**Figure 1D**). If VP^Gad1^ neurons contribute to relapse, then renewal should be reduced by their chemogenetic inhibition. We used a low dose of clozapine (0.1 mg/kg, i.p.) as the hM4Di ligand (Gomez et al., 2017) and have shown previously that this does not affect relapse behaviors in non-DREADD expressing rats (Gibson et al., 2018). Systemic injection of clozapine reduced renewal in Gad1-Cre+ animals (Context × Nosepoke × Group interaction: F _(1,13)_ = 5.024, p = .043) (**Figure 1E**). VP-dependent relapse can also be provoked by contingent re-exposure to alcohol (reacquisition) (Khoo et al., 2015) and this underpins a rapid return to drinking in humans (Marlatt and Donovan, 2005). We assessed whether VP^Gad1^ neurons contribute to this second form of relapse by pre-treating the two groups with 0.1 mg/kg clozapine and re-training them in a single session of self-administration. However, VP^Gad1^ chemogenetic inhibition had no effect on reacquisition (Nosepoke × Group interaction F _(1,13)_ = 0.054, p = .820) (**Figure 1E**). There was also no effect of VP^Gad1^ chemogenetic inhibition on locomotor activity when assessed in a locomotor chamber (Gad1-Cre-: 4705 ± 799 cm; Gad1-Cre+: 4901 ± 318 cm [mean ± SEM]) (F _(1,13)_ = .012, p = .9144).

Then we expressed the inhibitory hM4Di designer receptor in VP^PV^ neurons (AAV5-hSyn-DIO-hM4Di-mCherry) of LE-Tg(Pvalb-iCre)2Ottc (PV-Cre+) (*n* = 7) or PV-Cre− (*n* = 7) rats (**Figure 1F**) and chemogenetically silenced these neurons via injection of low dose clozapine (0.1 mg/kg, i.p.) prior to test. Similar to VP^Gad1^ neurons, VP^PV^ chemogenetic inhibition significantly reduced relapse during renewal (Context × Nosepoke × Group interaction: F _(1,13)_ = 9.47, p = .0088) (**Figure 1G**). However, in contrast to VP^Gad1^, chemogenetic inhibition of VP^PV^ neurons also reduced relapse during reacquisition (Nosepoke × Group interaction: F _(1,13)_ 4.73, p = .0486) (**Figure 1G**). There was no effect of VP^PV^ chemogenetic inhibition on locomotor activity when assessed in a locomotor chamber (PV-Cre-: 4908 ± 496 cm; PV-Cre+: 6555 ± 777 cm [mean ± SEM]) (F _(1,13)_ 3.362, p = .0897).

These results show distinct roles for VP neuronal populations in relapse. PV and Gad1, but not vGlut2, neurons express relapse-associated changes in c-Fos mRNA. Consistent with these changes, chemogenetic inhibition of Gad1 and PV neurons reduces relapse during renewal. However, chemogenetic inhibition of PV neurons additionally reduces relapse during reacquisition.

### VP^GAD1^ neurons promote relapse via lateral hypothalamus

VP^Gad1^ neurons control relapse during renewal but not reacquisition. VP neurons have extensive projections, including to lateral hypothalamus (LH), lateral habenula, and ventral tegmental area (VTA). LH is a likely target for VP contributions to relapse because LH has been strongly implicated in renewal of alcohol-seeking after extinction or punishment (Marchant et al., 2009; 2014). We combined retrograde tracing using CTb with c-Fos immunohistochemistry to study whether VP neurons projecting to LH are recruited during renewal (**Figure 2A**). Rats in groups ABA (n = 5), ABB (n =6) and AB0 (n = 3) received application of CTb-488 or CTb-555 (counterbalanced) to LH (**Figure 2C**) then were trained to self-administer alcoholic beer and extinguished prior to test for expression of extinction (ABB), renewal (ABA), or not tested (AB0). On test there was renewal in group ABA (Context × Nosepoke × Group interaction: F _(1,9)_ = 92.089, p < .00001) (**Figure 2B**). Renewal was associated with c-Fos protein expression in VP (ABA > ABB=AB0, F_(1,11)_ = 13.763, p = .003444), including in VP neurons retrograde labelled from LH (F_(1,11)_ = 56.99, p = .000099) (**Figure 2E**, **Table 3**), showing recruitment of the VP→LH pathway during relapse. To assess the role of the VP→LH pathway in renewal, we expressed halorhodopsin (AAV5-hSyn-eNpHR3.0-eYFP) (n = 6) or enhanced yellow fluorescent protein (eYFP) (AAV5-hSyn-eYFP) in VP (n = 5) and implanted optical fibres above LH (**Figure 2F**). We inhibited the VP→LH pathway for the duration of the 30 min test. ABA renewal was prevented by VP→ LH pathway inhibition (Context × Nosepoke × Group interaction: F _(1,12)_ = 39.33, p = .000041). The following day we tested these animals for reacquisition and again we inhibited the VP→LH pathway. However, there was no effect on relapse during reacquisition (Group × Nosepoke interaction: F_(1,12)_ = 0.269, p = 0.613) (**Figure 2G**).

**Figure 2.**
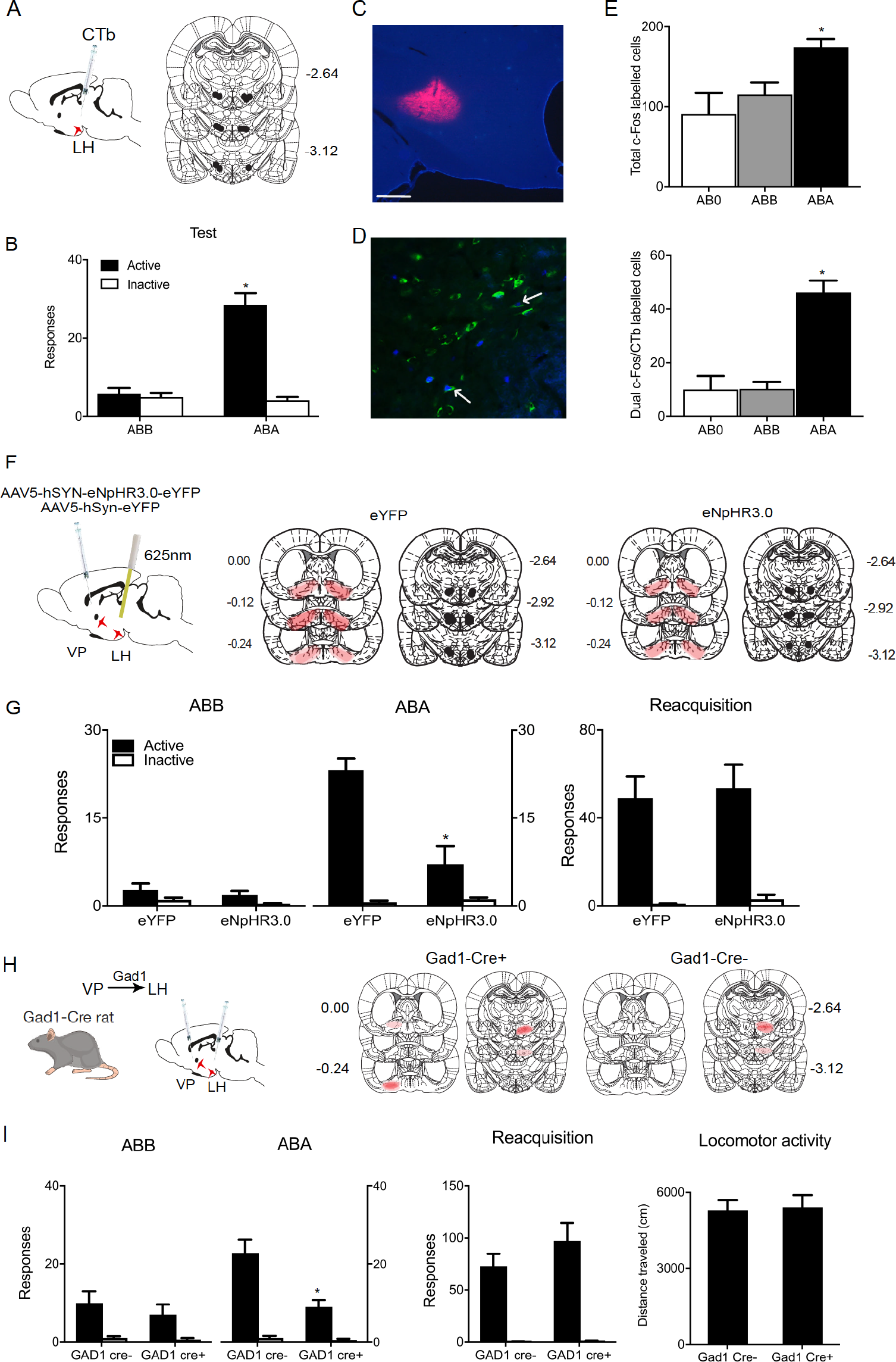
VP^Gad1^ neurons promote relapse via lateral hypothalamus. **(A)** and **(C)**, CTb-488 or CTb-555 was applied to lateral hypothalamus (LH) prior training and testing, scale bar = 200 μm. **(B)**, ABA renewal of alcohol seeking. **(D)**, Example of c-Fos (Blue), CTb (Green), and dual-labelling (shown by arrows) in VP. **(E)**, ABA renewal was associated with c-Fos expression in VP neurons including in VP→LH neurons. **(F)** Optogenetic inhibition of the VP→LH pathway and location of eNpHR3.0 and eYFP expression for all animals at 10% opacity; **(G)**, Optogenetic inhibition of VP → LH pathway reduced ABA renewal but not reacquisition **(H)**, Chemogenetic disconnection of VP Gad1 neurons from LH using contralateral disconnection with hM4Di in Gad1-Cre rats and location of hM4Di expression for all animals at 10% opacity. **(I)**, Chemogenetic disconnection of VP^Gad1^ → LH pathway reduced ABA renewal but reacquisition or locomotor activity.

These findings show that a VP→LH pathway is recruited to mediate relapse during renewal but not reacquisition. However, these findings do not identify the VP cell types responsible. VP^Gad1^ neurons project extensively to LH (Jennings et al., 2013) and the behavioral profile of VP→LH pathway inhibition resembled that of VP^Gad1^ inhibition, so we tested whether the role of VP^Gad1^ in renewal depends on LH. Using Gad1-cre+ (n = 5) and Gad1-Cre− (n = 5) rats we expressed the inhibitory hM4Di designer receptor in VP^Gad1^ neurons via AAV5-hSyn-DIO-hM4Di-mCherry in one hemisphere and the inhibitory hM4Di designer receptor non-selectively in the contralateral LH via AAV5-hSyn-hM4Di-mCherry (**Figure 2H**) to disrupt VP^Gad1^ and LH communication in Gad1-Cre+, but not Gad1-Cre−, rats. Renewal was reduced in Gad1-Cre+ rats (F _(1,8)_ = 5.35, p = .0493). (**Figure 2I**). Importantly, this same disconnection had no effect on relapse during reacquisition (F_(1,8)_ = 1.29, p = .2873) or on locomotor activity (F_(1,8)_ = .393, p = .5481) (**Figure 2I**). These effects of disconnecting VP^Gad1^ from LH recapitulated the effects on relapse of silencing VP^Gad1^ neurons, showing that the role of VP^Gad1^ neurons in relapse depends on LH.

### Convergence and segregation of ventral striatopallidal inputs to lateral hypothalamus

LH comprises a diversity of cell types relevant to appetitive motivation (Bonnavion et al., 2016). In mice, there are dense VP GABA inputs to LH GABA neurons (Jennings et al., 2013). We tested whether VP^Gad1^ neurons also provide inhibitory projections onto rat LH GABA neurons. To do this, we expressed ChR2 in VP^Gad1^ using AAV5-hSyn-DIO-ChR2(H134R)-eYFP and made whole-cell patch-clamp recordings from eYFP-labelled LH^Gad1^ neurons in Gad1-Cre+ rats (**Figure 3A**). Photostimulation (470 nm) of VP terminals evoked a short latency inhibitory responses (3.3 ± 0.8 ms from light onset; mean ± SD) in eYFP+ (i.e. LH^Gad1^) (53%, 10/19) but not eYFP-neurons (**Figure 3B, C**). The current’s kinetics, reversal potential, and sensitivity to the GABA-A receptor antagonist picrotoxin (100 μM; n = 4; **Figure 3D, E**) indicated that VP^Gad1^ neurons provide monosynaptic GABAergic inputs onto LH^Gad1^ neurons.

**Figure 3.**
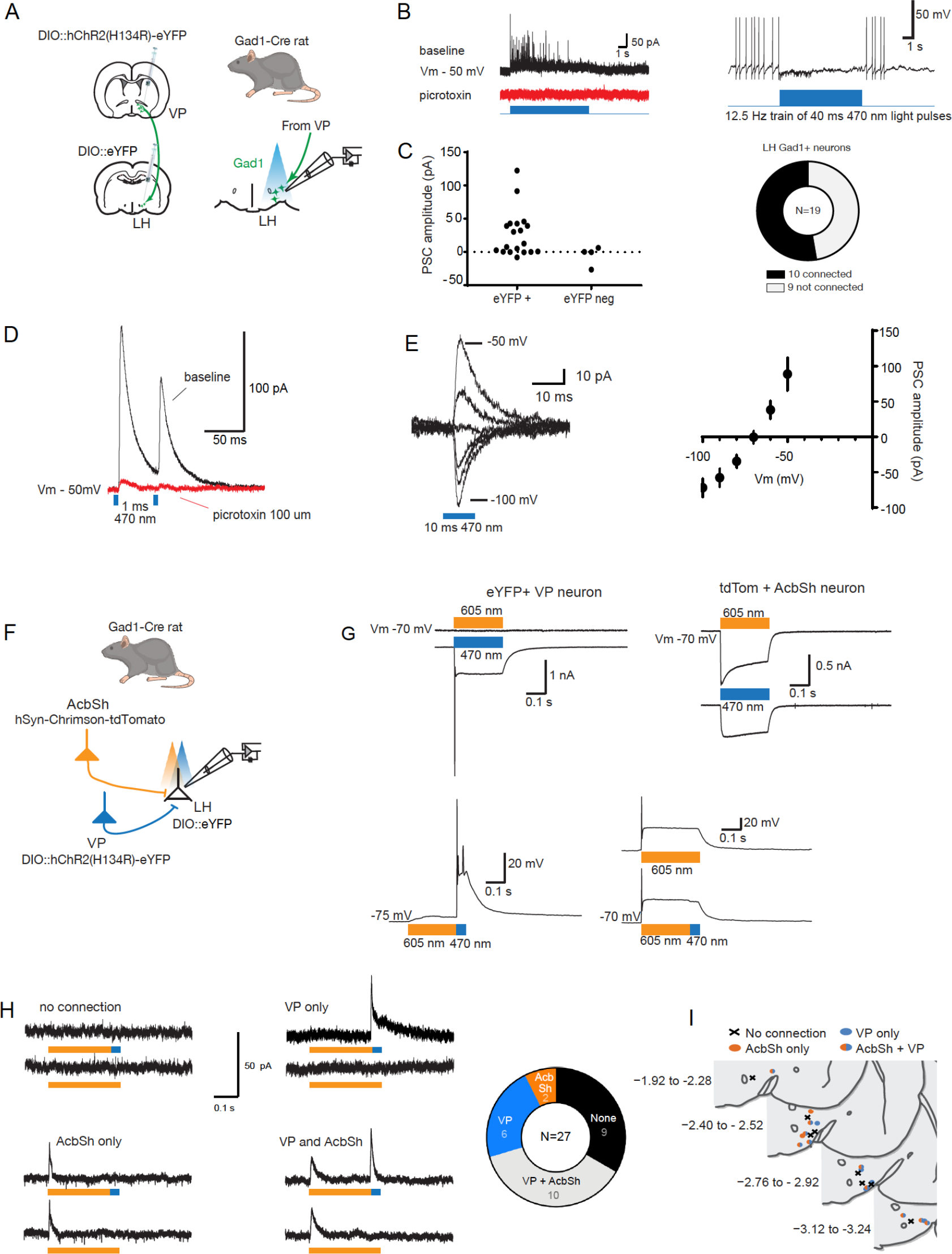
VP^GAD1^ neurons provide monosynaptic inhibitory GABAergic input to LHGad1 neurons. (A), Schematic for ChR2 mapping of VP inputs to LHGad1 neurons. (B) Representative responses of connected LHGad1 neurons to light pulse trains (40 ms, 470 nm at 12.5 Hz). Trains evoked fast picrotoxin-sensitive IPSCs. GABA-B receptor mediated slow IPSCs were not observed. (C), Amplitude of the light-evoked (470 nm, 10 ms) post-synaptic current (PSC; − 50 mV) in eYFP+ (LHGad1) and eYFP-neurons, with connectivity chart. Light evoked a reliable current in 10 of 19 (53%) LHGad1 neurons. (D), The light-evoked current (Vm – 50 mV) was blocked (96 ± 3%; Mean ± SD; n = 4) by bath application of the GABA-A receptor blocker picrotoxin (100 μM). (E), Individual example and summary data (mean ± SEM; n = 10) of the PSC evoked at holding potentials −100 to – 50 mV. The PSC reversal potential matches the Cl-equilibrium potential. (F), Schematic for testing convergence of VP and AcbSh inputs to LHGad1 neurons. (G) Spectral properties of Chrimson and ChR2 transduced neurons. ChR2 expressing VP neurons responded selectively to 470 nm (blue). Currents (Vm – 70 mV) evoked by blue and orange light are plotted above the voltage response evoked by an orange (250 ms)-blue (25 ms) light. Crimson expressing AcbSh neurons responded to 605 nm and 470 nm stimulation. Currents (Vm – 70 mV) evoked by blue and orange light are plotted above voltage responses to orange and orange-blue light. (H), Representative currents evoked in LHGad1 neurons by stimulation of AcbSh and VP terminals and connectivity chart. AcbSh inputs converged with VP inputs to LHG12ad1 neurons but VP inputs also provided additional segregated inputs to LHGad1 neurons. (I), Locations of LHGad1 neurons recorded as defined by source of input.

LH^Gad1^ neurons also receive monosynaptic GABA inputs from AcbSh (O'Connor et al., 2015; Gibson et al., 2018). Whereas VP GABA inputs to LH neurons promote relapse, AcbSh inputs prevent relapse (Gibson et al., 2018). These opposing roles in relapse raise the question of whether these two components of the ventral striatopallidal system target the same or different LH^Gad1^ neurons. To address this, we recorded from eYFP-labeled LH^Gad1^ neurons in Gad1-cre rats after expressing ChR2 in VP^Gad1^ neurons (AAV5-hSyn-DIO-ChR2(H134R)-eYFP) and the red-shifted opsin Chrimson (AAV5-hSyn-Chrimson-tdTomato) in AcbSh neurons (**Figure 3F**). This allowed us to identify AcbSh and VP inputs to the same LH^Gad1^ neurons via 605 nm and 470 nm optical stimulation respectively (**Figure 3G**). Using this approach, we could reliably optically-evoke currents in the majority of eYFP-labelled LH^Gad1^ neurons inputs (**Figure 3H**). There was convergence of ventral striatopallidal inputs, with 37% of LH^Gad1^ neurons receiving monosynaptic input from both AcbSh and VP. However, although 22% of LH^Gad1^ neurons responded selectively to VP^Gad1^ stimulation, only a minority (7%) responded selectively to AcbSh stimulation (**Figure 3H**). Thus, whereas AcbSh inputs converge with VP^Gad1^ inputs onto LH^Gad1^ neurons, there is further segregation of VP^Gad1^ inputs in the LH. There was no obvious relationship between ventral striatopallidal input selectivity and location of these Gad1 neurons in LH (**Figure 3I**).

### VP^PV^ neurons promote relapse via ventral tegmental area

VP^PV^ neurons control both renewal and reacquisition. So, we examined whether the role for VP^PV^ neurons in relapse also depends on LH. We used a three-group chemogenetic approach disconnecting VP^PV^ from LH. We expressed the inhibitory hM4Di designer receptor in VP^PV^ neurons via AAV5-hSyn-DIO-hM4Di-mCherry in one hemisphere and in either the contralateral (PV-Cre+ Contra n = 7) or ipsilateral (PV-Cre+ Ipsi n = 6) LH via AAV5-hSyn-hM4Di-mCherry (**Figure 4A, B**). This approach disconnects VP^PV^ from LH in group PV-Cre+ Contra but not PV-Cre+ Ipsi. In control PV-Cre− rats, we applied the hSyn-DIO-hM4Di to one VP and the hSyn-hM4Di to the contralateral LH (PV-Cre− Contra) (n = 7). We trained, extinguished, and tested these groups for relapse via renewal and reacquisition. Rats were injected with clozapine (0.1 mg/kg, i.p.) prior to both tests. In both renewal and reacquisition tests there was relapse (Renewal Context × Nosepoke interaction: F_(1,17)_ = 78.12, p < .00001; Reacquisition main effect of Nosepoke: F_(1,17)_ = 58.27, p < .00001) (**Figure 4C**) but there was no effect of chemogenetic VP − LH disconnection (Renewal Context × Group × Nosepoke interaction: F_(1,17)_ = 1.43, p = .481; Reacquisition Group × Nosepoke interaction: F_(1,17)_ = 1.81, p = .1961). There was also no effect of chemogenetic disconnection on locomotor activity (F_(1,17)_ = .274, p = .6074) (**Figure 4C**). So, the role of VP^PV^ neurons in relapse is independent of LH and anatomically dissociable from VP^Gad1^ neurons.

**Figure 4.**
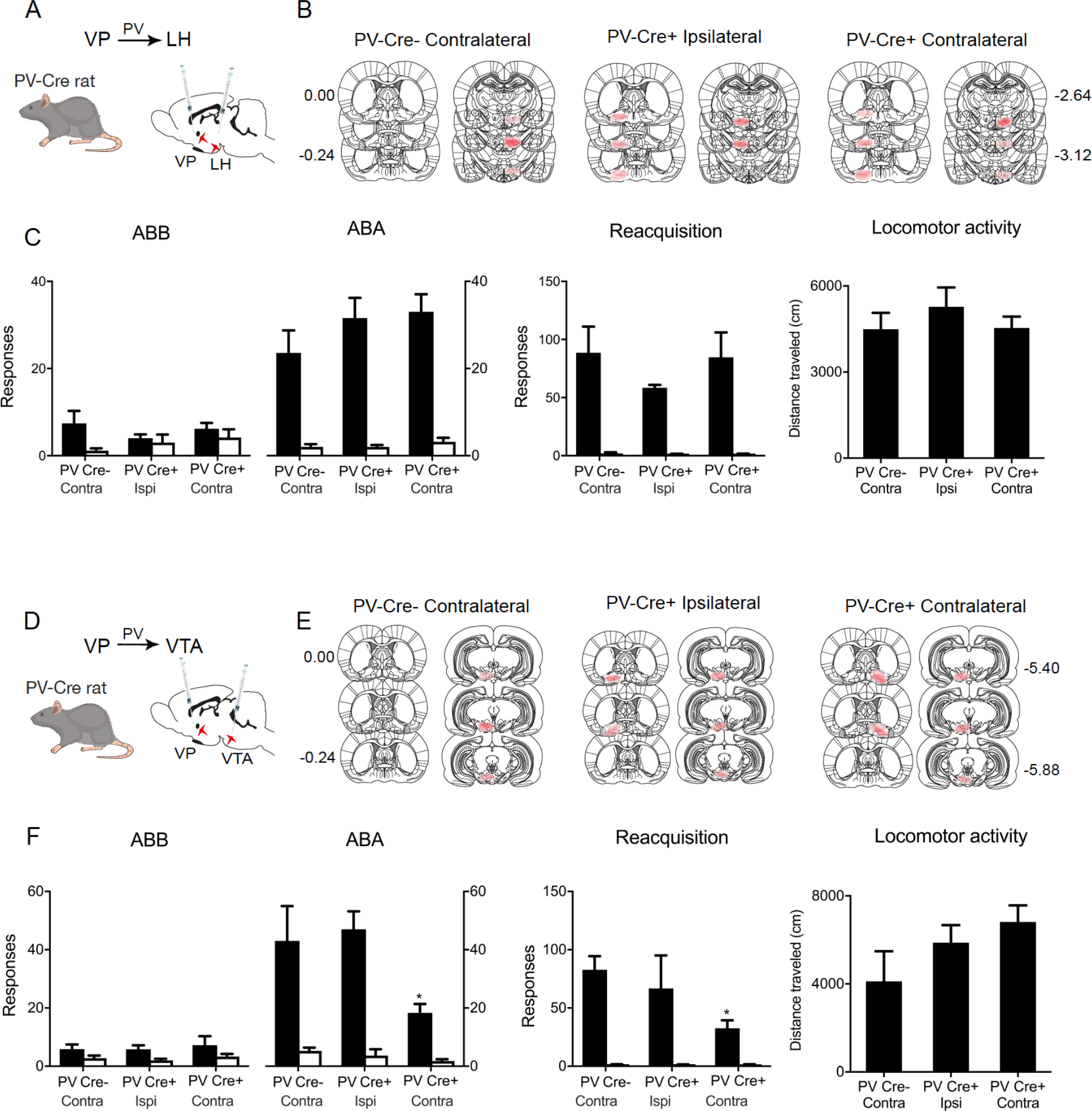
VP^PV^ neurons promote relapse via ventral tegmental area. **(A)** Cre-dependent inhibitory hM4Di was applied to VP of PV-Cre rats and hM4Di applied to LH of the same animals. **(B)**, Locations of hM4Di expression shown for all animals at 10% opacity. **(C)**, Chemogenetic disconnection of VP^PV^ → LH pathway had no effect on renewal, reacquisition, or locomotor activity. **(D)**, Cre-dependent inhibitory hM4Di was applied to VP of PV-Cre rats and hM4Di also applied to VTA of the same animals. **(E)**, Locations of hM4Di expression shown for all animals at 10% opacity. **(F)**, Chemogenetic disconnection of VP^PV^ → VTA pathway reduced renewal and reacquisition but not locomotor activity.

A VP →VTA pathway is a second plausible candidate for VP^PV^ contributions to relapse. VP^PV^ provide monosynaptic input to VTA dopamine and GABAergic neurons (Knowland et al., 2017; Faget et al., 2018). VP inputs to VTA are recruited during different forms of relapse (Mahler and Aston-Jones, 2012; Mahler et al., 2014; Prasad and McNally, 2016), and non-selective chemogenetic disconnection of VP neurons from VTA reduces relapse (Mahler et al., 2014; Prasad and McNally, 2016) To determine whether VP^PV^ contributions to relapse depend on VTA, we used the three group chemogenetic disconnection design (PV-Cre+ Contra n = 9, PV-Cre− Contra n = 7, PV-Cre+ Ipsi n = 6) (**Figure 4D, E**). We trained and extinguished rats prior to testing for relapse via renewal and reacquisition. Rats were injected with clozapine (0.1 mg/kg, i.p.) prior to tests. During both tests there was relapse (Renewal Context × Nosepoke interaction: F_(1,19)_ = 45.60, p < .00001; Reacquisition main effect of Nosepoke: F_(1,19)_ = 44.71, p < .00001) (**Figure 4F**). However, unlike VP^PV^ − LH disconnection, both forms of relapse were reduced by VP^PV^ − VTA disconnection (Renewal Context × Group × Nosepoke interaction: F_(1,19)_ = 8.13, p = .0011; Reacquisition Group × Nosepoke interaction: F_(1,19)_ = 5.49, p = .0315) (**Figure 4F**). There was no effect of this disconnection manipulation on locomotor activity (F_(1,19)_ = 2.67, p = .1206) (**Figure 4F**). Thus, chemogenetically disconnecting the VP^PV^ – VTA recapitulated the effects on relapse of silencing VP^PV^ neurons.

## Discussion

We studied the cellular and circuit architecture for relapse control in ventral striatopallidal pathways. VP has been viewed as an obligatory node in a ‘final common pathway’ for relapse. Different forms of relapse (cue, stress, prime, context) have different subcortical substrates, but these converge in dorsomedial prefrontal cortex to be routed through the ventral striatopallidal system generating relapse behavior (Kalivas and Volkow, 2005; Kalivas and O’Brien, 2007). This view has proved enormously useful in understanding the role of VP in relapse. Although our findings confirm this key role for VP, they show that this role is more complex still and that different forms of relapse are assembled from different VP neuronal populations and their projections.

### VP cell types in relapse

VP^Gad1^ neurons control relapse during renewal but not reacquisition. This depends, in part, on projections to LH where VP^Gad1^ neurons provide inhibitory monosynaptic GABAergic input to LH^Gad1^ neurons. So, the relapse phenotype observed after silencing VP^Gad1^ neurons was also observed after silencing VP^Gad1^ − LH interactions. AcbSh inputs to LH^Gad1^ neurons have the opposite role in relapse. Whereas VP^Gad1^ inputs to LH promote relapse, AcbSh GABAergic inputs to LH^Gad1^ neurons promote extinction of drug seeking and inhibit relapse (Gibson et al., 2018) as well as inhibit appetitive behavior more generally (O’Connor et al., 2015). Because LH^Gad1^ neurons are essential to encoding and using appetitive associations to guide behavior (Nieh et al., 2016; Sharpe et al., 2017), one possibility is that this convergence of AcbSh→LH^Gad1^ and VP→LH^Gad1^ pathways onto the same LH^Gad1^ neurons contributes to the balance between contextual control over extinction versus relapse. In this way, LH^Gad1^ neurons may serve a critical integrative role in promoting versus preventing relapse. Indeed, we identified considerable convergence between VP and AcbSh synaptic inputs to the same LH^Gad1^ neurons that could underpin such integration. However, there was also selective segregation of VP inputs from AcbSh inputs. That is, whereas AcbSh inputs tended to converge with VP inputs, VP inputs were additionally segregated from AcbSh inputs to LH^Gad1^ neurons. Whether these VP inputs represent parallel, independent channels of VP-derived information flow to LH^Gad1^ neurons or whether these inputs interact during relapse and other forms of appetitive motivation awaits investigation. Regardless, taken together with the literature, our results show opposing behavioral functions for VP and AcbSh GABAergic inputs to LH.

In contrast, VP^PV^ neurons control relapse during both renewal and reacquisition. This role for VP^PV^ neurons is independent of LH and depends on VTA. That is, the relapse phenotype observed after silencing VP^PV^ neurons was not observed after silencing VP^PV^ − LH interactions but was recapitulated after silencing VP^PV^ − VTA interactions. VP^PV^ neurons provide monosynaptic inputs to VTA GABA as well as dopamine neurons (Knowland et al., 2017) and both of these VTA cell types are important for relapse (Mahler et al., 2014; Gibson et al., 2018). However, the nature of VP input to these cell types differs. VP^PV^ input to VTA GABAergic neurons is inhibitory whereas VP^PV^ input to VTA dopamine neurons is predominantly excitatory (Knowland et al., 2017). So, different VP inputs to GABA and dopamine neurons could be linked to specific forms of relapse, but this awaits further investigation. Regardless, given the considerable overlap between VP^Gad1^ and VP^PV^ neurons, the differences we described here between these two cell populations was surprising and show functional as well as anatomical dissociations between the roles of these VP cell types in relapse.

Finally, there was no evidence here for recruitment of VP^vGlut2^ neurons selectively during relapse. Rather, VP^vGlut2^ neurons were recruited during test in either the extinction context (where relapse did not occur) or the training context (where relapse did occur). The role of VP^vGlut2^ neurons in promoting or preventing relapse remains to be determined. These neurons have been implicated in aversion and behavioral avoidance via their projections to the lateral habenula (Knowland et al., 2017; Faget et al., 2018) and this same projection acts to constrain reward seeking (Tooley et al., 2018). One common feature of tests in both the extinction and training contexts here was that the reinforcer was absent and these tests were under extinction conditions. So, recruitment of VP^vGlut2^ neurons during testing here may track this absence of the reward and constrain reward seeking accordingly.

### Methodological considerations

Our conclusion that VP^Gad1^ and VP^PV^ neurons serve complementary roles in relapse rests on multiple lines of evidence. Regardless, at least four methodological issues are worth considering. First, consistent with its location in the ventral basal ganglia, VP has a critical role in motor function and locomotor activity (Austin and Kalivas, 1990). This raises the possibility that the effects on relapse reported here were secondary to changes in locomotor activity. However, there was no evidence that relapse prevention was secondary to any locomotor deficits because locomotor activity was not altered under the conditions tested here. Second, consistent with recent findings (Gomez et al., 2017), we used low dose clozapine as the hM4Di actuator. This raises concerns about specificity of our chemogenetic manipulations. However, neither hM4Di expression nor clozapine injection alone affected relapse behavior (eg, **Figure 4c**). Rather, the impact of chemogenetic manipulations was selective and depended not just on the behavior measured but also the specific cell-type and region in which hM4Di was expressed. The same low dose clozapine injection in hM4Di expressing animals had no effect on relapse, caused a reduction of renewal but not reacquisition, or caused a reduction of both renewal and reacquisition, depending on the cells and pathways studied. Third, in one experiment we use relatively prolonged stimulation to optogenetically silence the VP→LH pathway during renewal. It is worth noting that this same stimulation had no effect in the same animals during test for reacquisition, so we consider it unlikely that the attenuation of renewal was due to non-specific effects. Regardless, our conclusion that VP → LH interactions mediate relapse during renewal was independently supported by both retrograde tracing and chemogenetic disconnection. Finally, consistent with our past work (Hamlin et al., 2007; Prasad and McNally, 2016; Gibson et al., 2018), rats had access to food and water for 1 hr per day. Under these conditions, rats could be responding for alcohol-rewarding effects, caloric effects, or a combination thereof. This is an important constraint on the generality of our findings. The relevance of the mechanisms we described here to other forms of relapse (cue, stress, priming reinstatement) to seeking other rewards (cocaine, heroin sucrose etc) awaits investigation.

## Conclusions

We studied the cellular and circuit architecture for relapse control in ventral striatopallidal pathways, showing how different forms of relapse are assembled from VP cell populations and their projections. Our findings add to an emerging literature showing that the functional diversity in the ventral striatopallidal system can be resolved by analyses of different families of VP neurons and their projection targets (Richard et al., 2016; Knowland et al., 2017; Faget et al., 2018; Tooley et al., 2018). In the context of relapse to drug seeking, our findings raise the possibility that treatments targeting these different ventral striatopallidal pathways may provide one approach to tailored interventions for different forms of relapse.

Paste Results here, use “merge formats” then select it all and click on the “Normal” style. Use “Heading 2” style for sub headings, “Heading 3” as needed for sub-sub headings

## Author Contributions and Notes

Data reported in the paper are archived in the UNSW Long Term Data Archive (ID: D0235271). This work was supported by the National Health and Medical Research Council (GNT1098436, GNT1138062), the Australian Research Council (DE170100509), UNSW Major Research Equipment and Infrastructure Initiative, and UNSW School of Psychology. We declare no competing interests.

Author contributions. <u>Conceptualization</u>: G.P.M. A.A.P., and J.P. <u>Experimentation</u>: A.A.P., C.X., C.C., and J.M.P. <u>Formal Analysis</u>: A.A.P., J.P., and G.P.M. <u>Writing – original draft</u>: G.P.M., A.A.P., and J.M.P. and <u>Writing – Review & Editing</u>: All authors. <u>Funding acquisition</u>: G.P.M., S.K., J.M.P., and A.A.P.

